# Subunit shuffling dynamics in KaiC’s central hub reveal the synchronization mechanism of the cyanobacterial circadian clock

**DOI:** 10.1101/2025.03.17.643614

**Authors:** Tian Yang, Zhuangcheng Zhen, Yuhai Tu, Qi Ouyang, Yuansheng Cao

## Abstract

Protein complexes are critical for cellular functions, and subunit exchange within these complexes is increasingly recognized as a key regulatory mechanism. In the cyanobacterial circadian clock, subunits shuffling of the core clock protein KaiC is thought to synchronize the clock, though the underlying mechanism remains unclear. We developed a chromatography-based method to monitor the shuffling dynamics of hexamerization domain of KaiC (KaiC-CI) and found that ATPase activity is essential for this process. By analyzing experiment data with quantitative models, we found that KaiC-CI hexamer stochastically disassembles into two oligomers for shuffling after hydrolysis. Further, by assuming a hidden conformation for post-hydrolysis hexamers, we established an ATPase activity-dependent model that quantitatively describes the shuffling dynamics of KaiC-CI hexamers, linking the shuffling rate to ATP hydrolysis and nucleotide exchange rates. Using this model, we estimated the shuffling dynamics of full-length KaiC with indirect experimental data. Our findings suggest that KaiC’s phosphorylation states regulate nucleotide exchange rates in the CI domain, thereby modulating ATPase activity and influencing subunit shuffling. This study provides a mechanistic framework for understanding the role of ATPase activity in subunit exchange and its implications for circadian clock regulation.

## Introduction

An emerging topic in the study of protein complexes is the functional role of their subunit exchange behavior^1^. Increasing numbers of experiments reveal that the subunit within protein complex can exchange within physiological timescales, potentially influencing the function of the complex. For example, in the study of the CaMKII holoenzyme, Stratton et al. and Bhattacharyya et al. found that when CaMKII are activated by Ca2^+^/CaM, the activation signal spreads through subunit exchange with non-activated proteins, implying that the exchange process may enhance neural signal transduction in synapse^2,3^. Similarly, in the study of the hexameric AAA+ ATPase protein P97, Huang et al. found that the subunit exchanging of P97 could be relevant to its substrate processing functions^4^. Another example is the study of the transcriptional factor NF-κB family, where homodimers of its core components can exchange subunits to form heterodimers, potentially regulating transcription^5^. These examples highlight an additional layer of protein-protein interaction by exchanging subunits to regulate their cellular functions. However, a quantitative understanding of subunit exchange, particularly activation-dependent exchange, is still lacking.

In the cyanobacterium, three proteins, KaiA, KaiB, and KaiC, interact to generate the circadian rhythm, where the phosphorylation level of the hexameric AAA+ ATPase KaiC oscillates stably both *in vivo* and *in vitro*^6-9^. In a noisy natural environment, the phosphorylation status of individual KaiC hexamer is heterogeneous. Several studies have proposed that monomer shuffling can synchronize the their phosphorylation levels, contributing to the robustness of the molecular clock^10-13^. Although early experiments indicated that KaiC hexamers exchange subunits during the dephosphorylation phase, molecular details of this shuffling mechanism remain unclear^14-16^. Quantitative models show that shuffling between KaiC hexamers consumes energy, which may be provided by ATP hydrolysis due to the ATPase activity in the CI domain ^10^. However, the relationship between ATPase activity and shuffling is not straightforward: the KaiC shuffling rate is higher during the subjective nighttime, when ATPase activity is relatively low, whereas shuffling rarely occurs when ATPase activity is high^14,17^. This apparent paradox highlights the need for detailed studies to elucidate the role of ATPase activity in subunit shuffling.

Generally, subunit exchange requires the dissociation of a subunit from the protein complex, which is often the rate-limiting step. KaiC consists of two domains: the CI domain, which alone forms a stable hexamer, and the CII domain, which exists in equilibrium between monomeric and hexameric states depending on its phosphorylation state^18,19^. These observations suggest that the CI domain is primarily responsible for hexamer assembly, while the stability of the hexamer is modulated by the phosphorylation state of the CII domain^20^. However, the exact influence of KaiC’s phosphorylation states on the hexamerization domain remains unclear. Although recent studies have shown the allosteric communication between the CII and CI domain, this static structural information is insufficient to fully explain the complex’s dynamic behaviors^21,22^.In this study, we first investigate the shuffling behavior of KaiC-CI hexamers and the role of ATPase activity in regulating shuffling dynamics. We then discuss how phosphorylation states influence subunit shuffling in full-length KaiC.

To quantify the shuffling dynamics of KaiC, we developed a chromatography-based approach to measure the distribution of hetero-hexamers formed by mixing native and epitope-tagged homo-hexamer proteins. First, we conducted a series of experiments to examine the shuffling behavior of the KaiC-CI hexamer and found that the ATPase activity is essential for subunit shuffling, although the exact relationship between the two remained unclear. Second, chromatography data suggested that the CI hexamer randomly disassembles into two oligomers, which was driven by the stochastic ATP hydrolysis cycle within the hexamer. Based on these experimental observations, we developed a dynamic model to describe the shuffling behavior of the KaiC-CI hexamer. We found that the shuffling rate is directly controlled by the ATP hydrolysis rate and nucleotide exchange rate. These two rates also determine the ATPase activity. This model quantitatively explains the complex relationship between shuffling rate and ATPase activity observed in experiments. Finally, although the model was initially proposed for the KaiC-CI hexamer, we demonstrated its applicability to full-length KaiC, providing insights into the allosteric regulation between the CI and CII domains.

## Results

### High-resolution chromatography reveals a strong dependence of subunit shuffling on ATPase activity

To monitor the subunit shuffling dynamics of the KaiC hexamers, we developed a method based on ion exchange chromatography (Fig. 1 *A*). Briefly, a 2×FLAG peptide was fused to the C-terminal of the protein, resulting in samples containing hexamers with varying amounts of 2×FLAG peptide. Due to the additional negative charges carried by the tag, hexamers with different tag quantities are eluted at distinct salt concentrations^23^. Then, the percentage of seven types of hexamers was recovered by fitting the raw signal with a combination of seven peak functions (method).

**Figure 1.**
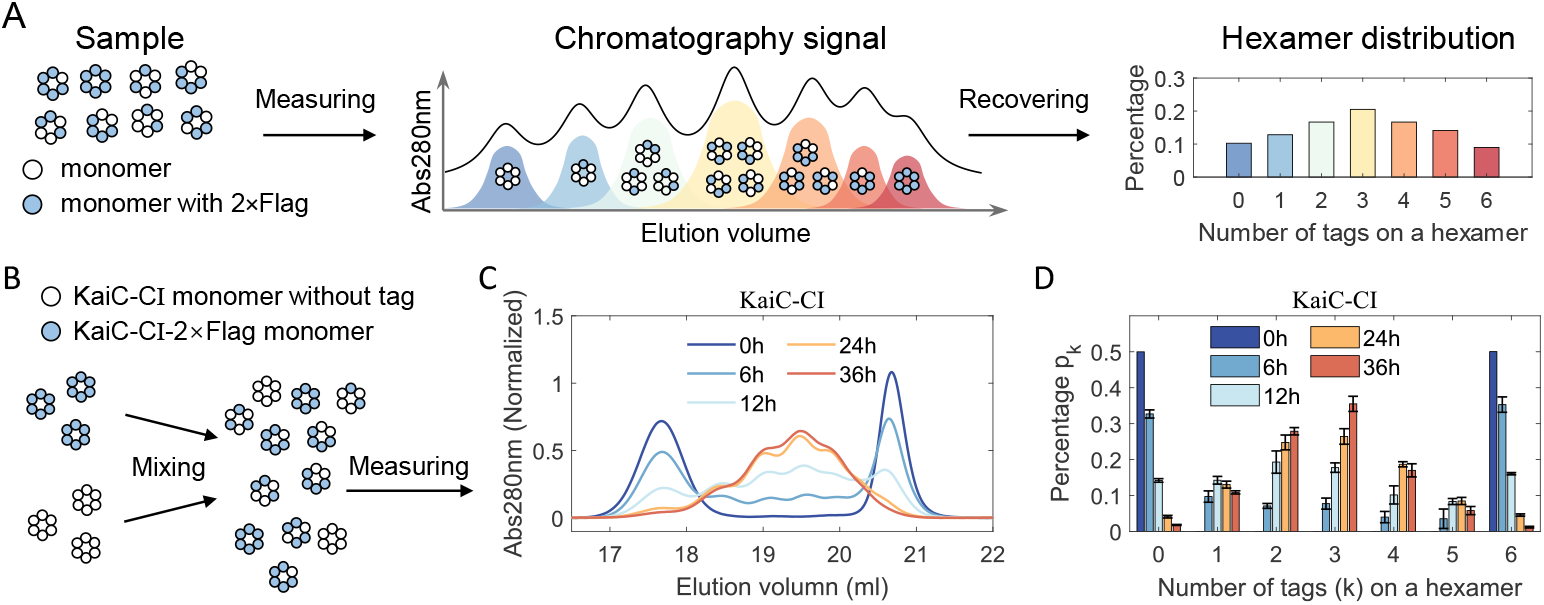
Measuring the compositional distribution of the KaiC hetero-hexamers by ion-exchange chromatography. (A)Schematic pipeline for measuring the hetero-hexamer distribution. Hexamer samples containing different numbers of 2×FLAG tags are separated by ion-exchange chromatography, and the hexamer distribution is recovered from the chromatography signal by fitting with a combination of seven peak functions. (B)Schematic of sample preparation and measurement for KaiC-CI. Native hexamers and 2×FLAG-tagged hexamers are mixed with a ratio of 1:1 in the reaction buffer. Then, samples are collected at different time points for chromatography analysis. (C)Chromatography signals for KaiC-CI hexamers. Peaks corresponding to homo-hexamers are centered for homo-hexamers are around 17.5ml and 20.5ml, respectively. Hetero-hexamers are eluted between 18ml and 20.5ml. (D)Recovered distribution for KaiC-CI hexamers from (C).

First, freshly purified native and 2×FLAG-tagged CI hexamers were mixed with a ratio of 1:1 and incubated in reaction buffer at 30°C (method). The mixed samples were analyzed by ion exchange chromatography at multiple time points (Fig.1 *B-D*). Initially, only two peaks corresponding to homo-hexamer were observed. After 6 hours of incubation, signals for hetero-hexamers were detected roughly between elution volumes of 18.5 ml and 20.5 ml, and the percentage of hetero-hexamers increased over the incubation time (Fig. 1 *C* and *D*, Fig. S1). We define the rate of hetero-hexamers formation as shuffling rate *k*_*s*_, which can be fitted with a one-phase exponential decay model (Fig. S2 and method). For wild-type CI, the shuffling rate is *k*_*s*_ = 0.091 ± 0.011*h*^−1^.

Second, to assess the effect of ATPase activity on shuffling, we studied CI hexamer mutants with varying ATPase activities^24^ (Fig. 2 *A*, Fig. S3-S4). The low ATPase activity mutant CI-S48T (0.09 *h*^−1^) hexamer exhibited a slow shuffling rate of 0.015 ± 0.004*h*^−1^. The CI-S157C mutant, with ATPase activity approximately two thirds of the wild-type (0.29 *h*^−1^ vs 0.43 *h*^−1^), displayed a similar shuffling trend to that of the wild type. In contrast, the high ATPase mutant CI-S157P (0.73 *h*^−1^) has approximately doubled ATPase activity compared with wildtype, while its shuffling rate is increased approximately 10-fold to 0.881 ± 0.030*h*^−1^. Noticeably, the hetero-hexamer percentage of CI-S157P quickly reached saturation within 6 hours. These results suggest that the ATPase activity and shuffling rate are not directly associated, indicating that intermediate processes are required to explain the factors influencing the shuffling rate (Fig. 2 *B*).

**Figure 2.**
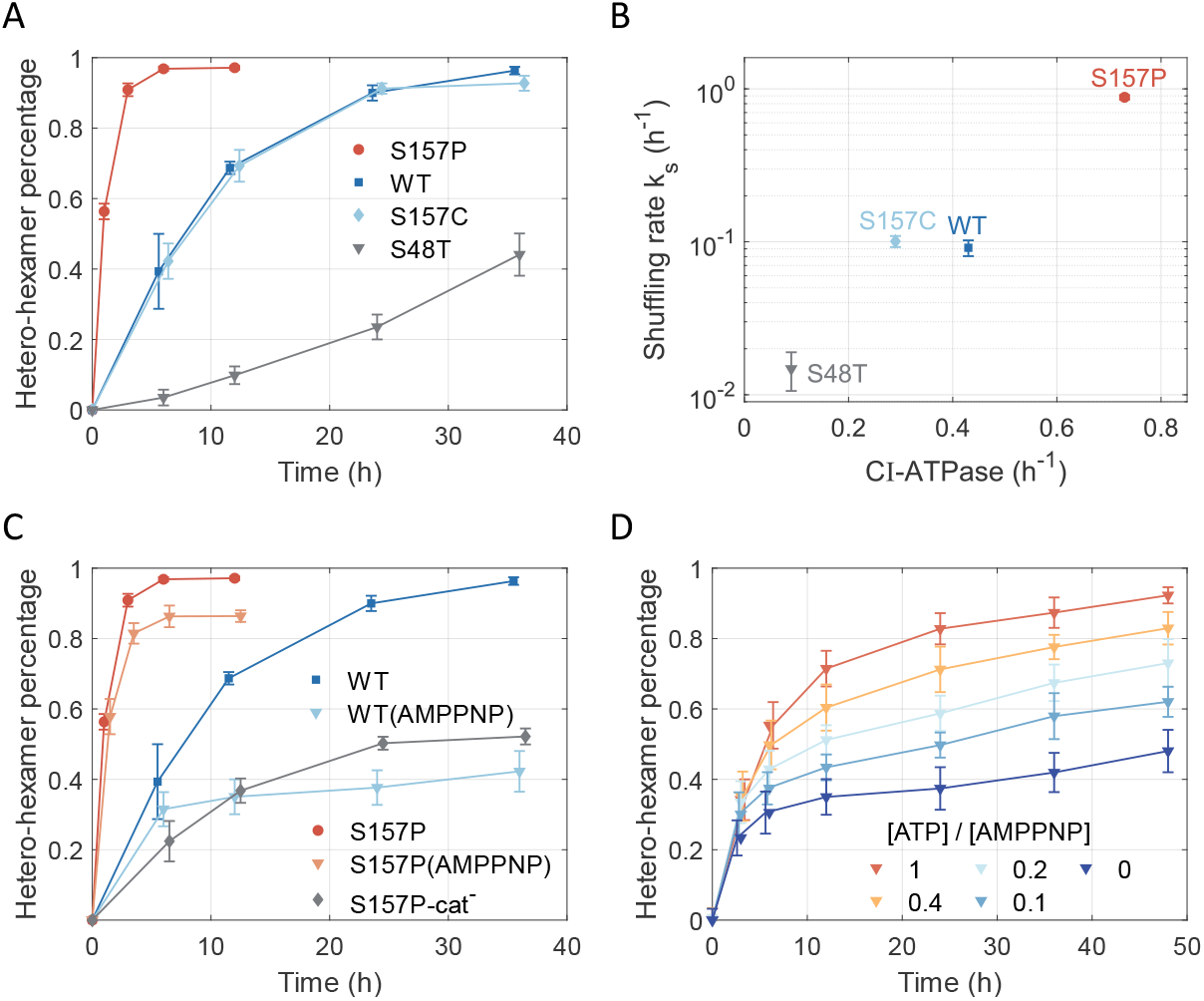
Shuffling dynamics of KaiC-CI hexamers. (A)Percentage of hetero-hexamers measured for KaiC-CI hexamers with different ATPase activity (two independent experiments; Fig. S1 and S3-S5). (B)Relation between shuffling rate and CI-ATPase activity. (C)Inhibition of shuffling dynamics by substituting ATP with AMP-PNP or by catalytic mutants. S157P-cat^-^ represents S157P/E77Q/E78Q mutant. (two independent experiments; Fig. S6-S7) (D)Percentage of hetero-hexamers at various ATP concentrations with 1mM AMP-PNP for KaiC-CI hexamers. (three independent experiments; Fig. S8-S10)

Next, we investigated the role of ATP hydrolysis in shuffling by substituting ATP with its non-hydrolyzable analog, AMP-PNP, in the reaction buffer. In the presence of AMP-PNP, the hetero-hexamer percentage at 36h decreased from 0.9 to 0.5 for wide-type CI (Fig. 2 *C*, Fig. S8 *A* and *B*), suggesting that ATP hydrolysis is essential for shuffling. We noticed that the fast-shuffling mutant CI-S157 still reached a high hetero-hexamer percentage under AMP-PNP condition, likely due to residual ATP bound to freshly purified proteins (Fig. 2 *C*, Fig. S6). To confirm this, we inhibited the ATPase activity of CI-S157P by mutating its catalytical site E77/E78 to E77Q/E78Q (denoted as CI-S157P-cat^-^). The shuffling behavior of this mutant closely resembled that of wide-type CI under AMP-PNP conditions (Fig. 2 *C*, Fig. S7).

Finally, we examined whether shuffling could be modulated by varying ratios of ATP and AMP-PNP concentrations. We measured the shuffling dynamics for wide-type CI hexamer in the presence of 1mM AMP-PNP and ATP concentrations ranging from 0mM to 1mM (Fig. 2 *D*, Fig. S8-S10). The percentage of hetero-hexamers decreased with lower ATP concentrations and higher AMP-PNP ratios. These results suggest that AMP-PNP-bound monomers are less likely to dissociate from hexamers, thereby reducing shuffling efficiency. Since nucleotide bind at the monomer-monomer interface, our findings imply that ATP binding stabilizes the monomer-monomer interface, consistent with previous experiments^25^. Notably, after the initial formation of hetero-hexamers within 12 hours, all conditions showed a slow and similar increase in hetero-hexamer percentage (Fig.2 *D*). At later stages, when AMP-PNP predominantly occupies nucleotide binding sites, hydrolysis becomes rare, resulting in slow shuffling.

### Spatially-independent ATPase activity drives random disassembly of KaiC-C I hexamers

As discussed above, inhibiting the ATPase activity of CI hexamer reduces its shuffling rate, indicating that the hexamer’s disassembly requires ATP hydrolysis. Previous molecular dynamics simulations suggest that KaiC-CII hexamer undergoes significant conformational change during nucleotide exchange^26^. Given the structure similarity between the CI hexamer and CII hexamer, it’s reasonable to propose that CI hexamer undergoes similar conformational change during nucleotide exchange^20^. Furthermore, several experimental studies have demonstrated that ATP hydrolysis in the CI domain induces conformational changes associated with KaiB binding^27-30^. These changes may also result in the disassembly of the CI hexamer and drive the observed shuffling behavior.

Crystal structures of various mutants of CI hexamers have shown that when ADP is bound at the monomer-monomer interface, the α6 and α7 helix in the counterclockwise monomer are repositioned^24^. This repositioning reduces the number of contacting residues at the interface (Fig. S11), weakening its stability. Thus, after ATP is hydrolyzed at one interface of the hexamer, that interface becomes less stable compared to the ATP-bound interface. We hypothesize that when two weakened interfaces coexist, the subunits between them are more likely to dissociate, resulting in hexamer disassembly (Fig. 3 *A*).

**Figure 3.**
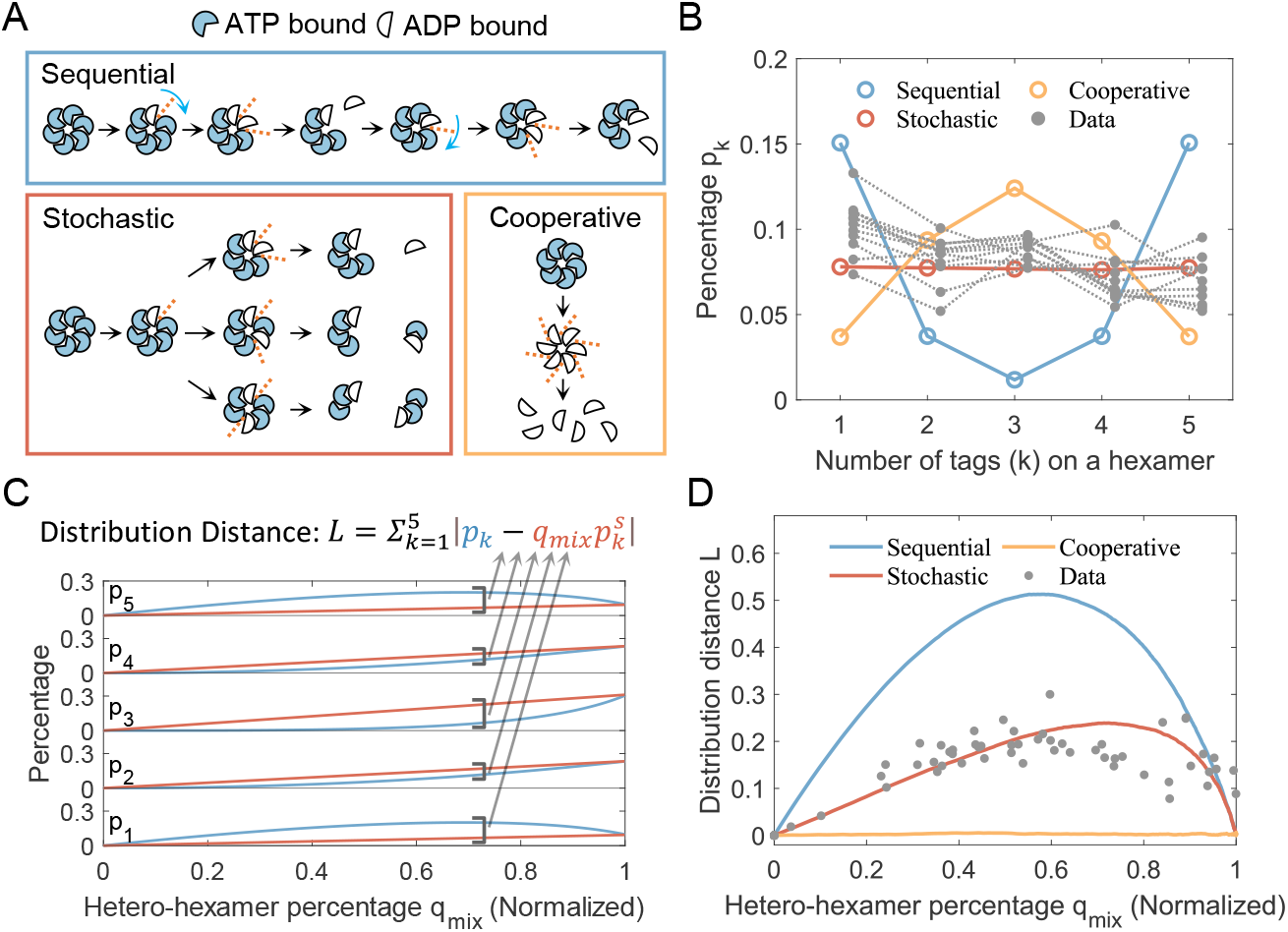
Inference of the dissociation pattern for KaiC-CI hexamers. (A)Schematic illustration of dissociation patterns corresponding to typical hydrolysis cycles. Dashed lines indicate unstable subunit-subunit interfaces. (B)Distribution of hetero-hexamers for three dissociation patterns when the percentage of hetero-hexamers reaches 40%. Data points are estimated by linear interpolation of all experimental measurements in this study (Fig. S1 and S3-S10). (C)Definition of the distribution distance *L*, calculated as the sum of the absolute errors relative to the normalized random distribution. (D)Comparison of the evolution trends of the distribution distance *L* between simulations and experimental data.

As an AAA+ ATPase, the KaiC-CI hexamer likely follows one of three common hydrolysis modes observed in other well-studied ATPase proteins^31^: sequential, cooperative, or stochastic. If the post-hydrolysis interface of a hexamer becomes loose and prone to breakage, then the hydrolysis pattern would primarily determine the dissociation process (Fig. 3 *A*). In the sequential hydrolysis mode, after ATP is hydrolyzed at the second interface, the hexamer disassembles into a monomer and a pentamer. In the cooperative hydrolysis mode, a hexamer dissembles into six monomers. In the stochastic hydrolysis modes, two loose interfaces are randomly generated, allowing equal production of subunits with varying numbers of monomers.

KaiC-CI alone forms stable hexamers, indicating that dissociation is the rate-limiting step of subunit exchange, as is generally the case in other protein complexes^32^. We performed stochastic simulations constrained by these three dissociation patterns (Fig. S12, methods). As shuffling progresses, the distributions of hexamers differ among the three modes. Fig. 3B illustrates the difference: when the hetero-hexamer percentage reaches about 40%, the sequential mode predominantly yields hetero-hexamers with 1 or 5 tags; the stochastic mode produces hetero-hexamers with nearly equal percentage; and the cooperative mode results in a random distribution of reconstructed hexamers. We estimate hetero-hexamer distribution when its percentage reaches 40% by linear fitting all experimental data. This analysis indicates that the dissociation mode of KaiC-CI is closest to stochastic (Fig. 3 *B*).

To further quantify the dissociation mode, we define a time-independent measurement *L* (Fig. 3 *C*), to assess the difference between these dissociation modes. The measurement *L* is expressed as

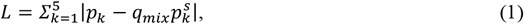

where *p*_*k*_ represents the observed distribution of hetero-hexamers,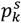 is the random distribution, and *q*_*mix*_ is the normalized percentage of hetero-hexamers. The random distribution and the normalized percentage are defined as

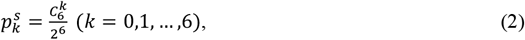

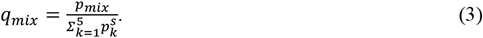

Here, *L* quantifies the distance between the observed distribution and random distributions at given mixing stage *q*_*mix*_ (Fig. 3 *D*). In sequential mode, the distance is always the highest among the three modes, while in cooperative modes, the distance is constantly zero. For stochastic mode, the evolution trend of distance *L* follows a trend similar to that of sequential mode, but the maximum value of *L* is reduced. Experimental data points closely align with the stochastic mode curve, implying that the CI hexamer undergoes random dissociation, with its hydrolysis pattern following a stochastic mode (Fig. 3 *D*).

### ATPase activity-dependent model explains the shuffling dynamics of KaiC-C I hexamers

Following the above discussion, we established a model to quantify the shuffling behavior of the CI hexamer. We assume that each monomer has two states: ATP-bound state (T state) and ADP-bound state (D state) (Fig. 4 *A*). Transitions between these states are described by the ATP hydrolysis rate *k*_*h*_ and nucleotide exchange rate *k*_*e*_(the reverse reactions of hydrolysis and nucleotide exchange are ignored but indicated in Fig. 4 *A*). Thus, the ATPase activity *k*_*cat*_ is defined as the net ATP consumption rate in one hydrolysis and exchange cycle at a nonequilibrium steady state

**Figure 4.**
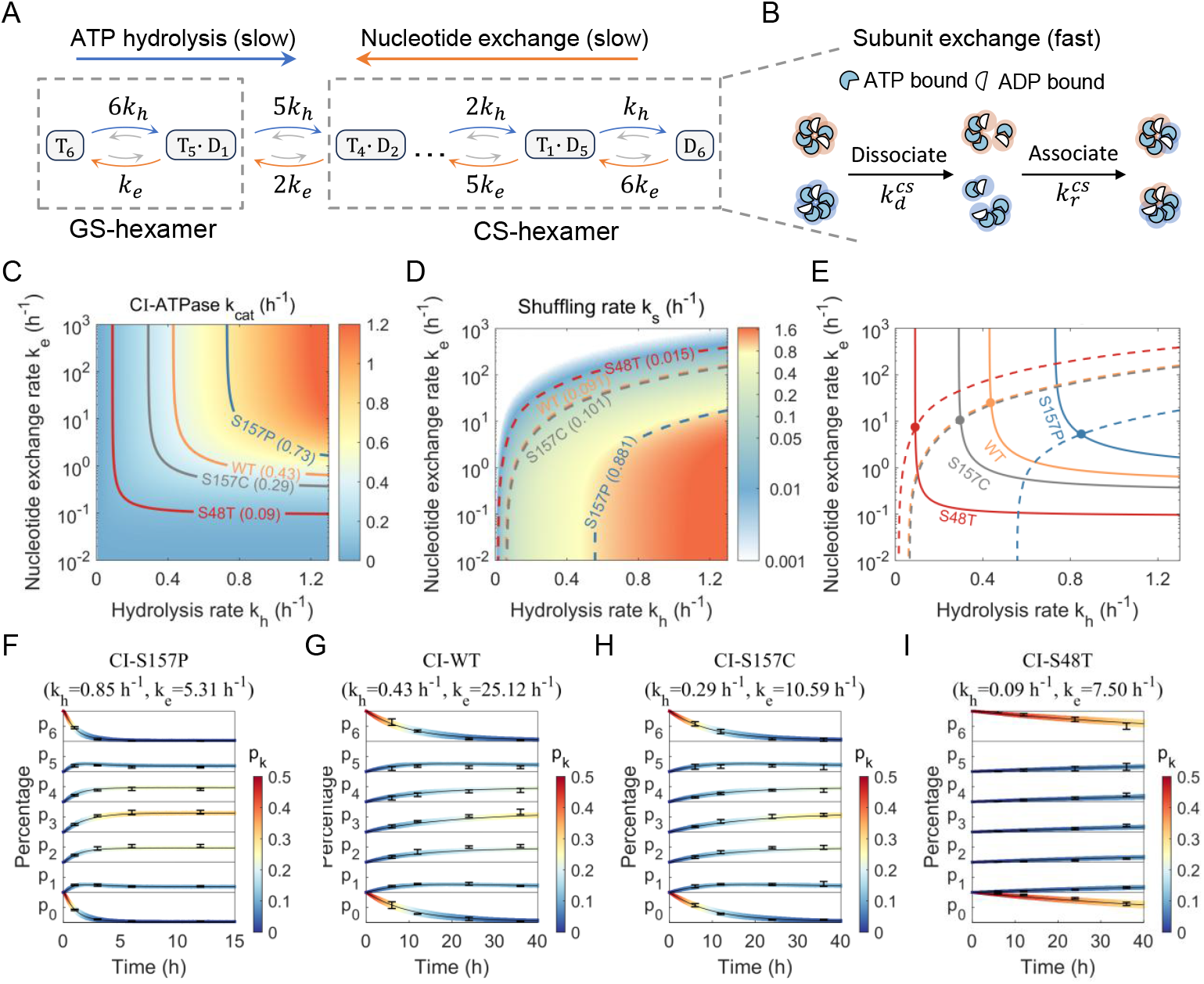
Model of subunit shuffling for KaiC-CI hexamers. (A)Schematic of ATPase activity-related reactions. T(D) represents a monomer bound with ATP(ADP). Hexamers with less than two ADP are ground-state hexamers (GS-hexamers) and with two or more are competent-state hexamers (CS-hexamers). (B)Schematic of the shuffling process for CS-hexamers. Subunits are traced using two background colors to distinguish their origins. (C-E) Fitting of model parameters (hydrolysis rate and nucleotide exchange rate) for KaiC-CI hexamers. Diagrams for CI-ATPase activity (Eq. 4) and shuffling rate *k*_*s*_ are shown. The optimal parameters are determined by the intersection of ATPase contour and shuffling rate contour. (F-I) Evolution of hetero-hexamer distribution fitted by the model. The width of the curves represents the fitting range (method), and black markers are experimental data.

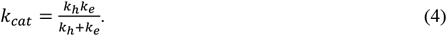

When a hexamer contains two or more ADP-bound monomers, it has a higher propensity to disassemble. As previously proposed^33^, we refer to such hexamers as the competent state hexamer (CS-hexamer). Conversely, a hexamer with fewer than two ADP-bound monomers is considered a ground-state hexamer (GS-hexamer). We assume that subunit exchange occurs among the CS-hexamers with a dissociation rate 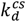 and an association rate 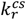 (Fig. 4 *B*). The model is governed by the master equation

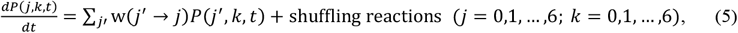

where *P*(*j, k, t*) represents the probability of the hexamer having j ATP-bound monomers and k tags at time *t. w* denotes the transition rate (*k*_*h*_ or *k*_*e*_) for ATPase activity-related reactions (Fig. 4 *A*). The shuffling reactions account for the disassembly and assembly reactions of CS-hexamers (Fig. 4 *B*). Disassembly occurs when there are two or more ADP-bound monomers (CS-hexamers) present. New hexamers form randomly with available oligomers.

The dissociation rate 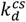 is estimated to be approximately 1 *s*^−1^, which is much faster than the hydrolysis and nucleotide exchange reactions (SI). This means the CS-hexamers’ subunits are well mixed before ATPase-related reactions occur. Thus, we assume the shuffling reactions reach fast equilibrium and model them with a stochastic algorithm (methods).

We solve the master equation numerically (methods). The evolution of distribution of the tag state is calculated as

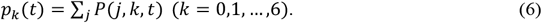

This time-dependent distribution is used to numerically fit the shuffling rate *k*_*s*_, which depends only on the hydrolysis rate *k*_*h*_ and the nucleotide exchange rate *k*_*e*_:

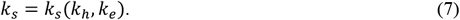

Simulation results show that the shuffling rate is higher in the low *k*_*e*_ region for a given ATPase activity (Fig. 4 *C* and *D*). In the high *k*_*e*_ region, the shuffling rate remains relatively low, even when the ATPase activity is high.

Eq.(4) and Eq.(7) are sufficient to determine *k*_*h*_ and *k*_*e*_ given the ATPase activity *k*_*cat*_ and shuffling rate *k*_*s*_ from the dynamic model in Eq. (6). We then fitted the model parameters using constraints from ATPase activity and experimentally measured shuffling rates (Fig. 4 *C-E*). The results demonstrate that the model accurately reproduces the shuffling dynamics of CI hexamers (Fig. 4 *F-I*).

Our results indicate that the ATPase activities of CI-S157P, CI-S157C, and CI-S48T hexamers are mainly affected by changes in their hydrolysis rate *k*_*h*_, while their nucleotide exchange rate *k*_*e*_ remains around 10 *h*^−1^(Fig. 4 *E* and Table S1). Since the percentage of ATP and ADP bound on a hexamer is determined by the hydrolysis and nucleotide exchange rate, our model also predicts the nucleotide-binding states of the CI hexamer, consistent with previous studies^25,34^ (SI and Fig. S13). Additionally, for experiments involving AMP-PNP to inhibit hydrolysis, we found the binding affinity of KaiC for ATP and AMP-PNP differs, and the affinity ratio *K*_*A*_ between ATP and AMP-PNP is in the range from 7.4 to 23.8 (Fig. S14 *A* and SI). For experiments carried out with AMP-PNP, the best fit for the nucleotide exchange rate *k*_*e*_ shifts from 25.5 *h*^−1^ in the ATP condition to 12.0 *h*^−1^ in the AMP-PNP condition (Fig. S15). This reduction in *k*_*e*_ is likely due to the lower total concentration of the ATP and AMP-PNP in the AMP-PNP contained buffer (Fig. S15 *G-I*).

### The implication of the model for full-length KaiC

We also applied our chromatography approach to monitor the shuffling dynamics of full-length KaiC. For the dephosphorylation mimic mutant KaiC-AA (S431A, T432A), subunit exchange was not observed using our method (Fig. S16 *A* and *B*), consistent with previous measurements^14,16^. For the phosphorylation mimic mutant KaiC-EE (S431E, T432E), our method failed to resolve its hetero-hexamer distribution due to the presence of non-hexamer oligomers, which eluted before the hexamer peak (Fig. S17 *A*). These oligomers may be dissociated from CS-hexamers, as their proportion decreases when ATP hydrolysis is inhibited^33^ (Fig. S17 *B* and *C*). Our model established for KaiC-CI hexamer also provides a quantitative relation for the fraction of CS-hexamer as (method)

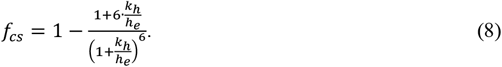

Since previous experiments suggest that CS-hexamers are dominant when S431 of KaiC is phosphorylated (32), we used this information to infer the model parameters of full-length KaiC.

For KaiC-AA, its CI ATPase activity is about 19.0 ATP/day/monomer (0.79 *h*^−1^) and almost no CS-hexamers are observed^33,35^. With these constraints, the shuffling rate is estimated to be less than 0.05*h*^−1^(method and Fig. 5 *A-C*). Similarly, the CI ATPase activity of KaiC-EE is about 2.0 ATP/day/monomer (0.08 *h*^−1^) and its fraction of CS-hexamer is evaluated around 95%^33,35^, resulting a shuffling rate of approximately 0.8*h*^−1^. In this scenario, the hydrolysis rate *k*_*h*_ of the two phosphor mimic mutants is similar, while their nucleotide exchange rate *k*_*e*_ in the CI domain varies significantly, from 10^3^*h*^−1^ to 10^−1^*h*^−1^.

**Figure 5.**
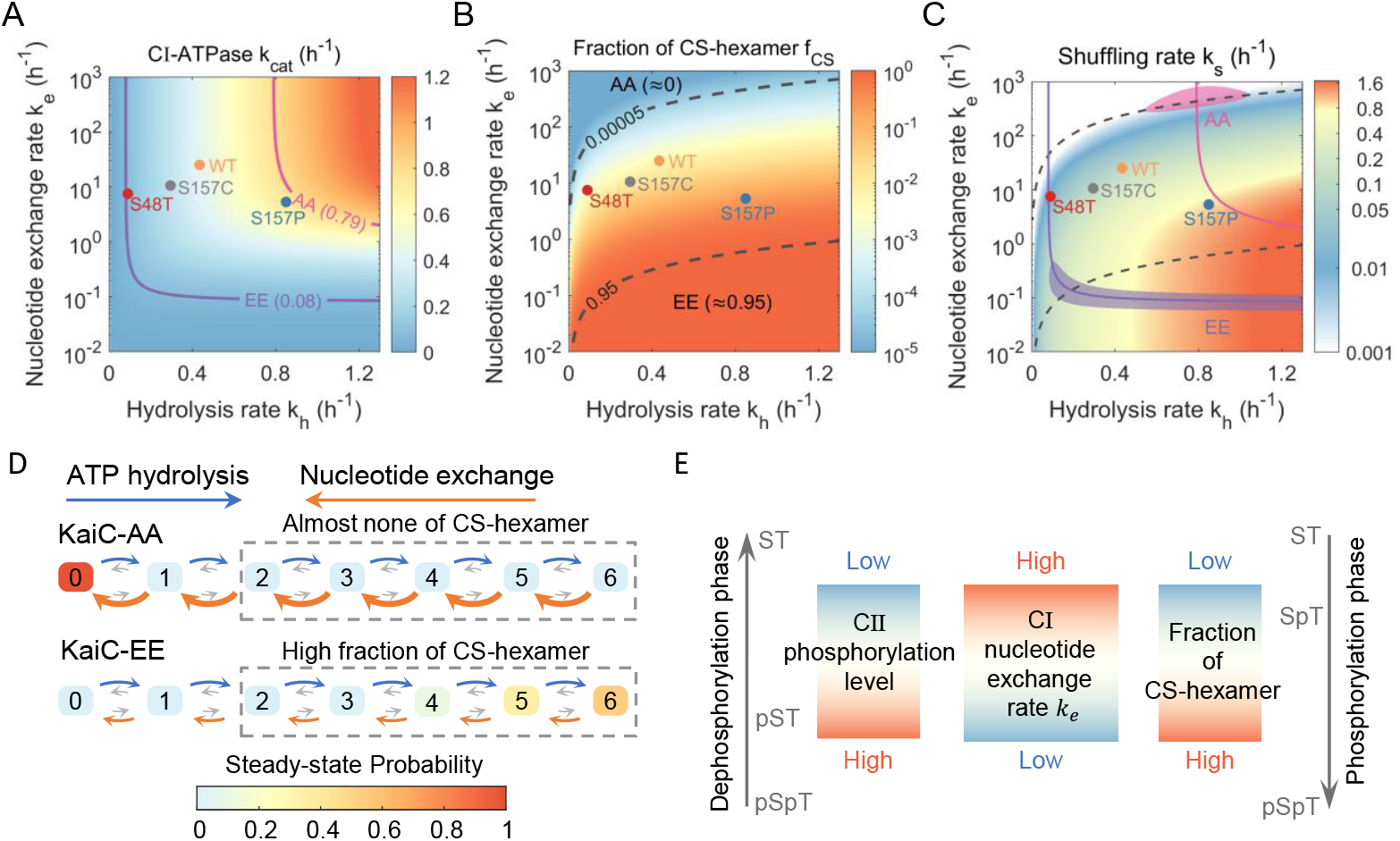
Effect of CII phosphorylation state on subunit shuffling. (A-C) Fitting of model parameters (hydrolysis rate and nucleotide exchange rate) and prediction of shuffling rate *k*_*s*_ for KaiC hexamers. Diagrams for CI-ATPase activity (Eq. 4), fraction of CS-hexamer, and shuffling rate *k*_*s*_ are shown. The parameter range for KaiC-AA and KaiC-EE are estimated based on their fraction of CS-hexamer and CI-ATPase activity (method). The optimal parameters of four KaiC-CI hexamers are also indicated. (D)Estimated Steady-state distribution of nucleotide states in the CI hexamers for KaiC-AA and KaiC-EE (method). The value in the box corresponds to the amount of ADP bound to CI hexamer, and the size of the arrow reflects the relative value of the reaction rates. (E)Schematic depiction of the regulation of subunit shuffling through the phosphorylation states of CII domain. Phosphorylation at S431 reduces the nucleotide exchange rate in the CI domain, leading to an increased proportion of CS-hexamers. These CS-hexamers represent the conformational state in which subunit shuffling occurs.

Our model explains the seemingly contradictory relationship between shuffling rate and ATPase activity in full-length KaiC. For KaiC-AA, despite its high CI ATPase activity, its nucleotide exchange rate is much higher than its hydrolysis rate, reducing the likelihood of GS-hexamers transitioning to CS-hexamers, thereby lowering dissociation efficiency (Fig. 5 *D*). Since the S157P mutant greatly increases the shuffling rate for KaiC-CI hexamer, we tested whether this effect could make KaiC-AA hexamer shuffle. However, no exchange behavior was observed for KaiC-AA-S157P hexamer within 16 hours (Fig. S16 *C*). This result aligns with our model, as the S157P mutant primarily increases *k*_*h*_ (ATP hydrolysis rate) but does not reduce *k*_*e*_ (nucleotide exchange rate), which is insufficient for KaiC-AA to enter the shuffling region (Fig. 5 *C*).

For KaiC-EE, the observable shuffling behavior is due to its nucleotide exchange rate being comparable to its hydrolysis rate, leading to a high fraction of CS-hexamer available for subunit shuffling, despite its low overall CI ATPase activity (Fig. 5 *D*). Similarly, introducing the S48T mutation to KaiC-EE maintained a high percentage of CS-hexamers, even though the shuffling rate of the CI-S48T hexamer was low (Fig. S17 *D*). This indicates that *k*_*e*_ of KaiC-EE is sufficiently low such that reducing *k*_*h*_ with S48T mutant does not shift it out of the shuffling region (Fig. 5 *C*).

The differences between KaiC-AA and KaiC-EE suggest that CI ATPase activity in full-length KaiC is mainly controlled by the nucleotide exchange rate *k*_*e*_. This rate is possibly modulated by the phosphorylation states of the CII domain, which in turn affects the shuffling behavior (Fig. 5 *E*).

## Discussion

In this paper, we incorporated quantitative models with biochemical measurements to elucidate the molecular mechanisms underlying subunit shuffling in KaiC hexamers, especially the role of the ATPase activity in this process. By inspecting the shuffling behavior of KaiC-CI hexamers with a chromatography method, we found that the ATPase activity is essential for subunit shuffling. However, the precise relationship between ATPase activity and shuffling could not be fully resolved through biochemical measurements alone. To bridge the gap between macroscopic experimental data and microscopic molecular mechanisms, we applied quantitative models for data analysis. Firstly, by examining the shuffling dynamics with a model only constrained by protein complex dissociation, we found that KaiC-CI hexamers randomly disassemble into two oligomers, corresponding to the stochastic hydrolysis pattern of ATPase protein. Secondly, stochastic simulations incorporating a transient conformational state (CS-hexamer), in which hexamers exchange subunits, demonstrated that our model explains the experimental data and provides a quantitative relation for the shuffling rate. In this model, ATPase activity is decomposed into two processes: ATPase hydrolysis and nucleotide exchange. The rates of these processes determine both ATPase activity and shuffling rate, thereby establishing a clear link between shuffling and ATPase activity.

Although our chromatography approach could not fully resolve the shuffling dynamics of full-length KaiC, our quantitative model enables us to infer these dynamics using indirect experimental observations. For full-length KaiC, model parameters were estimated based on the CI ATPase activity and the fraction of CS-hexamer, which also gave a consistent estimation for the shuffling rate. Furthermore, our approach clarified the inconsistency between the shuffling rate and the CI ATPase activity in full-length hexamer and provided insights into the allosteric communication between CI and CII domain of KaiC. Previous studies have suggested that the phosphorylation of S431 decreases the CI ATPase activity^17,35^. Our analysis shows that the CI ATPase is mainly tuned by changes in the nucleotide exchange rate, which in turn alters the fraction of CS-hexamers available for shuffling (Fig. 5 *D* and *E*).

Recent structural studies have revealed a relationship between the phosphorylation states of S431 and structural changes in the CI domain, showing that phosphorylation of S431 increases the preference for ADP in the CI nucleotide pocket^21^, which is consistent with our hypothesis. Although large conformational changes associated with hexamer disassembly have not been reported in current structural studies, we anticipate that future studies--such as cryo-electron microscopy(cryoEM)—may provide a clearer view of CS-hexamers, shedding light on the regulatory mechanism between CI and CII domain. Additionally, our quantitative model allows us to estimate the nucleotides distribution within a hexamer (Fig. S13), which could aid in analyzing experimental data, such as single-particle cryo-EM to reveal the conformational landscape of the protein complex at atomic resolution^36-39^.

Our quantitative model for subunit shuffling provides a foundation for more detailed studies on the functional role of subunit exchange in the circadian clock. Our results suggest that the hydrolysis pattern of CI hexamer is stochastic, which is quite different from previous models^10,40,41^. For full-length KaiC, further research is needed to determine the exact hydrolysis pattern of CI hexamer. However, since the CI ATPase activity is modulated by the phosphorylation level, the hydrolysis pattern of CI hexamer in full-length KaiC could be influenced by the phosphorylation cycle in CII domain, as well as the allosteric communication between the two domains^21,22,35,42^. This insight is key for detailed modeling of subunit shuffling in cyanobacterial circadian clock proteins. In our framework, subunit shuffling happens when the phosphorylation level is high (Fig. 5 *E*). This state of hexamer coincides with KaiB binding, suggesting that subunit dissociation may facilitate KaiB binding during the dephosphorylation phase of the clock, when the KaiB-binding segment in the CI domain is more frequently exposed ^28^. In this context, shuffling plays a role in synchronizing the clock at high phosphorylation levels. Conversely, when the phosphorylation level is low, shuffling occurs less frequently, and KaiA may play a role in synchronizing the clock into a low phosphorylation level^7,43-48^. Modeling the dynamics of shuffling in response to changing phosphorylation levels will help clarify the relationship between these regulatory elements.

In conclusion, our study presents a quantitative model for ATPase activity-related subunit shuffling, shedding light on the role of ATPase activity in the subunit shuffling of KaiC. Moreover, our model predicts a hidden conformation of KaiC protein and provides a potential feedback regulation mechanism mediated by phosphorylation signals, which will require further experimental validation. Our framework of subunit exchange is not limited to KaiC and can be applied to other activation-triggered subunit exchange processes, offering insights into the functional regulation in these molecular machines.

## Material and Methods

An extended SI Appendix, Material, and Methods section is provided and includes the following: Molecular cloning and protein purification; Sample preparation and chromatography analysis; Processing of the chromatography data; Stochastic simulation for subunit shuffling dynamics; Estimation for the dissociation constant for CS-hexamer; Stochastic simulation of ATPase related subunit exchange dynamics; Fitting of the experimental data; Steady-state distribution of the nucleotide states; Estimating parameters for KaiC-AA and KaiC-EE.

## Acknowledgments

The work by Q.O., T. Y. and Z. Z. is supported by the National Natural Science Foundation of China (12090054). The work by Y.C. is supported by the National Natural Science Foundation of China (2024YFA0919600). The work by Y.T. is supported by NIH grant (R35GM137134). This research is supported by High-performance Computing Platform of Peking University.

