## Supporting Information for "Subunit shuffling dynamics in KaiC’s central hub reveal the synchronization mechanism of the cyanobacterial circadian clock"

**This PDF file includes:**

Supporting text

Figures S1 to S17

Tables S1

SI References

### Material and methods

#### Molecular cloning and protein purification

Plasmids of wide-type KaiC were synthesized by Genscript on the pGEX-4T-1 vector. All mutants were generated using the Q5 site-directed mutagenesis kit (E0554S; New England Biolabs) and validated by sequencing. Hexamers of KaiC and KaiC-CI (1-250) were purified following a previous protocol[1]. Briefly, plasmids were transformed to BL21(DE3) competent cells. Proteins were expressed at 16°C overnight induced by 5μM IPTG; then, the bacterium was lysed by ultrasonication with an ice bath. Proteins were initially collected by GST beads (20562ES03; Yeasen), and the GST tags were cleavage on beads by thrombin (20402ES03; Yeasen) at 4°C overnight. After removing GST tags, proteins were purified by ion-exchange chromatography using a Hitrap-Q HP 1ml column (Cytiva) and size-exclusion chromatography using a Superdex 200 Increase 10/300 GL column (Cytiva). After purification, all proteins are stored at -80°C in a storage buffer (20mM Tris-pH8.0, 150mM NaCl, 10% Glycerol, and 1mM DTT).

#### Sample preparation and chromatography analysis

Native and 2×Flag tagged KaiC-CI hexamers are mixed in reaction buffer (20mM Tris-pH8.0, 150mM NaCl, 5mM ATP (A2383; Sigma-Aldrich), 5mM $MgCl_2$, 0.5mM EDTA) at 30°C with equal molar concentration and the final concentration is 10μM. For experiments substituting ATP with AMP-PNP (A2647; Sigma-Aldrich), the concentration of AMP-PNP is 1mM. Ion exchange chromatography buffer A contains 20mM Tris-pH8.0 and 10% Glycerol. Buffer B contains 20mM Tris-pH8.0, 1M NaCl and 10% Glycerol. 500μl samples were collected and loaded to a Mono Q 5/50 GL column (Cytiva) for analysis at 4°C. Samples are eluted applying a 0 to 900 mM NaCl linear gradient with a length of 40 column volume at a flow rate of 0.8 ml/min.

#### Processing of the chromatography data

The raw 280nm abortion signal of chromatography is fitted with a combination of seven Vogit functions as

$$raw\ signal = \sum_{k=0,1,\dots,6} A_k \cdot V(x - x_k; \beta_k; \gamma_k), \tag{1}$$

where $A_k$ is the amplitude of the peak for hexamers with $k$ tags, $x_k$ is the center of the signal, $\beta_k$ and $\gamma_k$ is the width parameter of the Voigt function. Then the distribution of hexamers with different tags are recovered as

$$p_k = \frac{\int A_k \cdot V(x - x_k; \beta_k; \gamma_k) dx}{\sum_k \int A_k \cdot V(x - x_k; \beta_k; \gamma_k) dx}\ (k = 0,1,\dots,6). \tag{2}$$

To fit the distribution, controlling experiments with only homo hexamers are firstly measured to determine the center $x_0$ and $x_6$, then the other five centers of hetero hexamer can be recognized directly. Then the peak parameter $A_k$, $\beta_k$ and $\gamma_k$ are fitted by nonlinear least-squares data-fitting method with predetermined peak centers. To estimate the fitting error, peak centers are randomly perturbed around the predetermined positions to repeat the fitting procedure. Then, the final distribution is recovered by calculating the average of ten fittings, and the fitting error is estimated as the standard deviation $\sigma_B$ of these fittings. For repeated experiments, the final error is

$$\sigma = \sqrt{\sigma_A^2 + \sigma_B^2}, \tag{3}$$

where $\sigma_A$ is the standard deviation for repeated experiments.

#### Stochastic simulation for subunit shuffling dynamics

First, we consider the subunit exchange dynamics between two types of homo-dimer $A_2$ and $B_2$ with reactions: $A_2 \leftrightarrow 2A,\ B_2 \leftrightarrow 2B,\ A + B \leftrightarrow AB$. The dissociation rate and association rate are denoted by $k_r$ and $k_d$ respectively. When the concentrations of both $A_2$ and $B_2$ are well above the dissociation constant $k_d/k_r$, it can be shown the rate of subunit exchange is limited by $k_d$ and the formation of hetero-dimer $AB$ can be expressed as

$$[\mathrm{AB}]_{(\mathrm{t})} = [\mathrm{AB}]_{(\mathrm{t}=0)}\mathrm{e}^{-\mathrm{k_d t}} + [\mathrm{AB}]_{(\mathrm{t}=\infty)}(1 - \mathrm{e}^{-\mathrm{k_d t}}). \tag{4}$$

Generally, when two types of homo-oligomer $A_n$ and $B_n$ are mixed, it's challenging to solve corresponding dynamic equations directly as for a dimeric protein. Assuming the subunit exchange dynamics are limited by the dissociation rate $k_d$, then the shuffling dynamics can be conveniently simulated by the Gillespie algorithm with τ-Leaping[2]. For the stochastic dissociation mode, each hexamer has a propensity of $k_d\tau$ to dissociate into two subunits during each time interval τ, and these dissociated subunits are randomly associated to form hexamers. For sequential mode, hexamers are only allowed to dissociate into a monomer and pentamer. the cooperative mode, hexamers are directly dissociated into six monomers (Fig. 3 *A*).

#### Estimation for the dissociation constant for CS-hexamer

Typically, the association rate $k_r \sim 10^7 M^{-1}s^{-1}$ for stable protein complex[3]. The concentration of KaiC in standard reaction condition is $3.5\mu M$, which implies the dissociation constant $K_D$ for CS-hexamers is about $0.1\mu M$. Then the dissociation rate is estimated as $k_d = k_r \cdot K_D \sim 1\ s^{-1}$, which is much faster than the ATPase related reactions.

#### Stochastic simulation of ATPase-related subunit exchange dynamics

Here, we extended dissociation-limited subunit shuffling dynamics to be limited by ATPase-related reactions. The nucleotide and tag states of hexamers are denoted by matrix $N$ and $T$, respectively (size of $N_{total} \times 6$). Then, the nucleotide state of a monomer is represented by $N_{i,j}$ with values 0, 1, and -1 for ATP bound, ADP bound, and AMP-PNP bound states, respectively. Similarly, $T_{i,j} = 1$ for a monomer with tag and $T_{i,j} = 0$ for a monomer without tag. For convenient, we define a function $\delta_k(N_{i,j}) = 1$ when $N_{i,j} = k$, then the propensities for all reactions are:

*Hydrolysis* $\quad w_{i,j}^{h} = k_h \cdot \delta_0(N_{i,j}),$ (5)

*Nucleotide exchange (ATP)* $\quad w_{i,j}^{b\to ATP} = P\{ATP\} \cdot k_e \cdot \delta_1(N_{i,j}),$ (6)

*Nucleotide exchange (AMPPNP)* $\quad w_{i,j}^{b\to AMPPNP} = P\{AMPPNP\} \cdot k_e \cdot \delta_{-1}(N_{i,j}),$ (7)

Where $P\{ATP\}$ and $P\{AMPPNP\}$ are determined as

$$\frac{P\{ATP\}}{P\{AMPPNP\}} = K_A \frac{[ATP]}{[AMPPNP]}. \tag{8}$$

$K_A$ is the ratio of the ATP's affinity for KaiC to the AMP-PNP's affinity for KaiC.

During a short time interval τ (much shorter than $k_h^{-1}$ or $k_e^{-1}$), the nucleotide states of hexamers are updated at first; after that, all hexamers with two or more ADP-bound monomers (CS-hexamers) are disassembled into two subunits according to their positions of ADP monomers and conjugated subunits were exchanged randomly to form new hexamers.

Additionally, a slow dissociation rate for hexamers occupied with AMP-PNP is required to fit the experimental data. Thus, we also introduce a reaction for the dissociation of hexamers with 1 or 0 ADP monomers (GS-hexamers) with a rate of $k_d^{gs}$ when AMP-PNP is supplied in the reaction buffer.

During the simulation, the tag distribution $p_k$ is updated as

$$p_k = \sum_{i=1}^{N_{total}} \delta_k\left(\sum_{j=1}^{6} T_{i,j}\right)/N_{total}. \tag{9}$$

Nucleotide distribution is updated similarly.

**Fitting of the experimental data**

For the sample reacting in ATP buffer, its hetero-hexamer percentage $p_{mix}$ is fitted by the one-phase exponential decay model as

$$p_{mix}(t) = p_{mix}(t = \infty) \cdot (1 - e^{-k_s t}), \tag{10}$$

Where $k_s$ is defined as shuffling rate (Fig. S2). Here, the pre-factor is given by random distribution (Eq. 3 in main article) as

$$p_{mix}(t = \infty) = \frac{\sum_1^5 p_k^s}{\sum_0^6 p_k^s} = \frac{2^6-2}{2^6}. \tag{11}$$

The relation $k_s = k_s(k_h, k_e)$ is determined by fitting equation (10) for data generated by stochastic simulation. The ATPase activity is expressed as

$$k_{cat} = \frac{k_h k_e}{k_h + k_e}. \tag{12}$$

Then the model parameters are constrained by

$$\theta^2(\hat{k}_h, \hat{k}_e) = \frac{(\hat{k}_s - k_s)^2}{k_s^2} + \frac{(\hat{k}_{cat} - k_{cat})^2}{k_{cat}^2}. \tag{13}$$

The optimal parameters are the intersection of two contours (Fig. 4 *C-E*). The parameter range is estimated within the region where $\theta^2 < 0.05$. 50 independent simulations with parameters randomly selected from the region are carried out, and all simulations are drawn to represent the fitting range.

For samples incubating with AMP-PNP, the data is directly fitted by stochastic simulation. First, the dissociation rate for GS hexamers is directly fitted as $k_d^{gs} \sim 0.017 h^{-1}$ with data at later time points, where most monomers are bounded with AMP-PNP, and the increasing trend of hetero-hexamer percentage is slow. Then the parameters $k_e$, $k_h$, and $K_A$ are fitted with constrain of ATPase activity, i.e., $k_{cat}$ is fixed as $0.43\, h^{-1}$ for KaiC-CI. The parameters are estimated with the R-square with weight as:

$$R^2 = 1 - \frac{\sum_i w_i \cdot (\hat{y}_i - y_i)^2}{\sum_i w_i \cdot (\bar{y}_i - y_i)^2}, \tag{14}$$

$$w_i = \frac{1}{\sigma_i^2}. \tag{15}$$

Where weight $w_i$'s are introduced to comprise the error of experimental data and $\sigma_i$ is the error calculated by equation (3). Additionally, to avoid overfitting, the $R^2$ is calculated with hetero-hexamer percentage (Fig. S14). 100 independent simulations with parameters randomly selected from the region where $R^2 > 0.95$ are carried out and all simulations are drawn to represent the fitting range.

**Steady-state distribution of the nucleotide states**

The dynamic equation of nucleotide states with n monomers in a complex is expressed as

$$\begin{cases} \frac{dq_0}{dt} = -nk_h q_0 + k_e q_1 \\ \frac{dq_k}{dt} = -[(n-k)k_h + kh_e]q_k + (n-k+1)k_h q_{k-1} + (n-k-1)k_e q_{k+1}\ (k = 2, \dots n-1) \\ \frac{dq_n}{dt} = -nk_e q_n + k_h q_{n-1} \end{cases} .$$

(16)

Where $q_k$'s are probability for a hexamer bound with k ADP (n=6 for hexamer). The steady-state distribution is solved as

$$q_k = \frac{C_n^k\left(\frac{k_h}{k_e}\right)^k}{\left(1+\frac{k_h}{h_e}\right)^n}. \tag{17}$$

**Estimating parameters for KaiC-AA and KaiC-EE**

The fraction of CS-hexamers is expressed as

$$f_{cs} = \sum_{k=2}^{6} q_k = 1 - \frac{1+6\cdot\frac{k_h}{h_e}}{\left(1+\frac{k_h}{h_e}\right)^6}, \tag{18}$$

Then the model parameters are constrained by

$$\theta^2(\hat{k}_h, \hat{k}_e) = 0.2 \cdot \frac{\left(\hat{f}_{cs}-f_{cs}\right)^2}{f_{cs}^2} + \frac{\left(\hat{k}_{cat}-k_{cat}\right)^2}{k_{cat}^2}. \tag{19}$$

We added a weight in the first term to comprise the uncertainty for the fraction of CS-hexamers. CI-ATPase activity and fraction of CS-hexamer are adapted from[4,5]. For KaiC-AA, $f_{cs} = 5 \times 10^{-5}$ and $k_{cat} = 0.79\,h^{-1}$. For KaiC-EE, $f_{cs} = 0.95$ and $k_{cat} = 0.08\,h^{-1}$. The parameter range is estimated within the region where $\theta^2 < 0.1$ (Fig. 5 *A*-*C*).

Fig. 5 *D* is plotted according to the model's estimation, with hydrolysis rates for KaiC-AA and KaiC-EE both at $0.8h^{-1}$, and nucleotide exchange rate of $500h^{-1}$for KaiC-AA and $0.1h^{-1}$ for KaiC-EE.

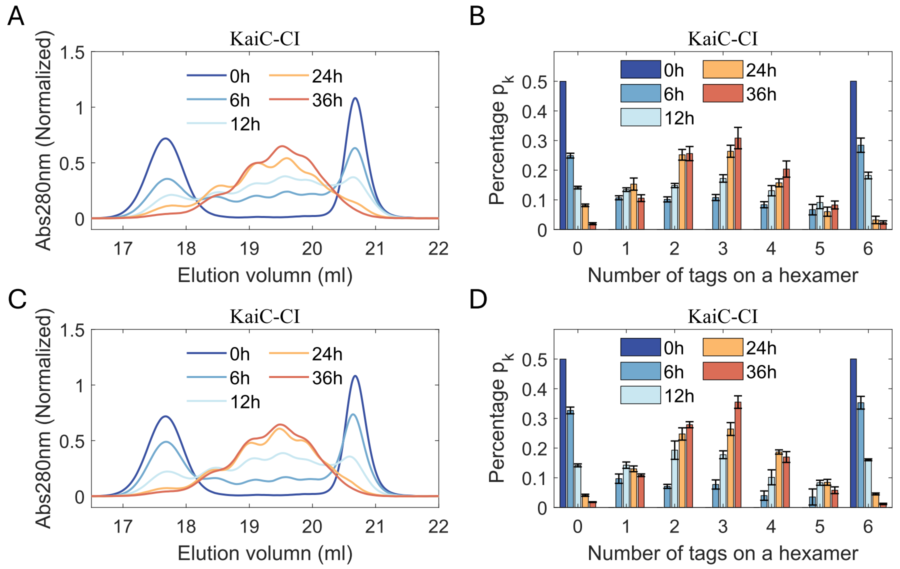


**Fig. S1.** Chromotography signals and recovered distributions for two independent experiments that measure the shuffling dynamics of KaiC-CI hexamers (C-D) are also shown in Fig. 1.

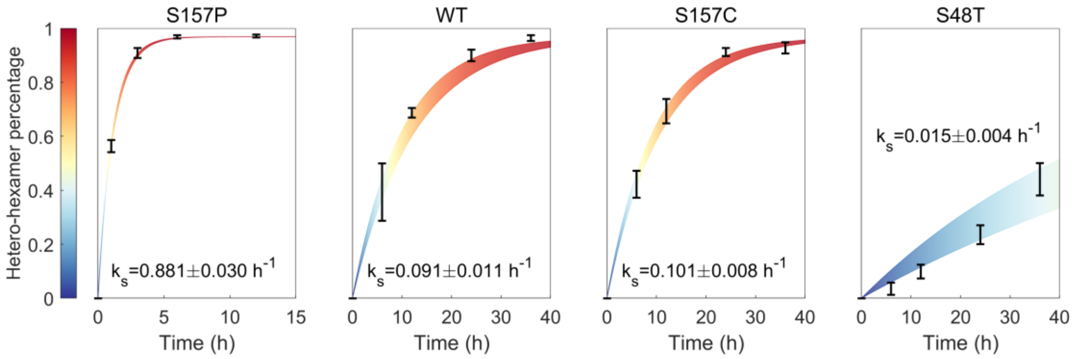


**Fig. S2.** Fitting for sample incubated in 5mM ATP buffer. The hetero-hexamer percentage is directly fitted by shuffling rate $k_s$.

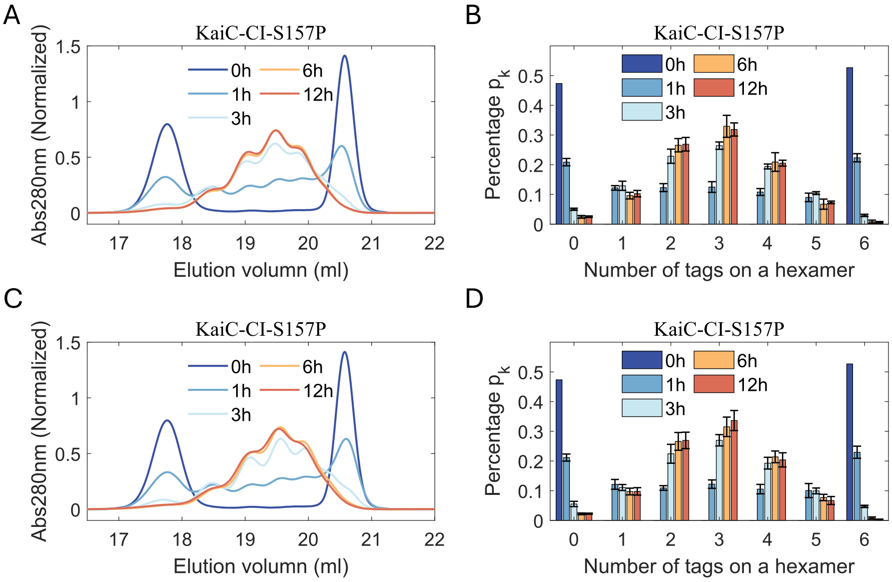


**Fig. S3.** Chromotography signals and recovered distributions for two independent experiments that measure the shuffling dynamics of KaiC-CI-S157P hexamers.

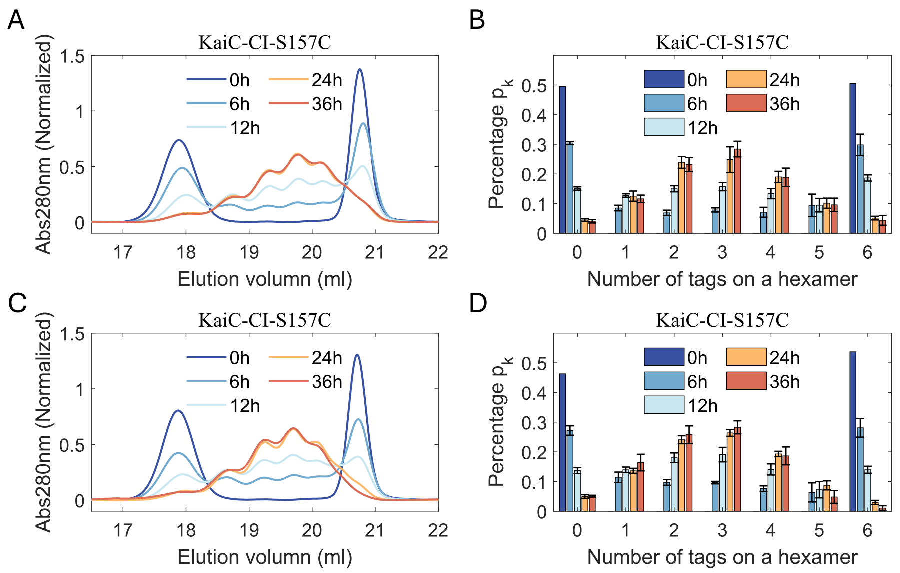


**Fig. S4.** Chromotography signals and recovered distributions for two independent experiments that measure the shuffling dynamics of KaiC-CI-S157C hexamers.

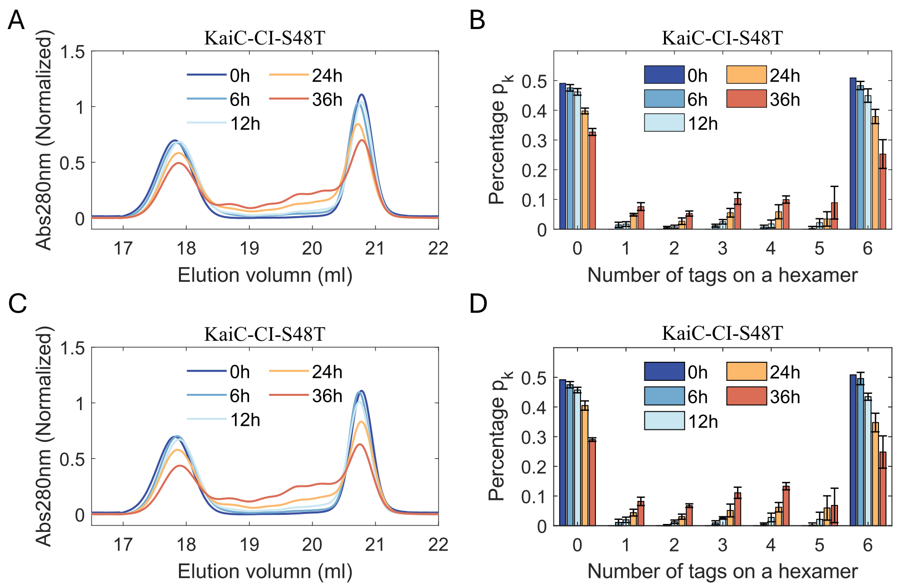


**Fig. S5.** Chromotography signals and recovered distributions for two independent experiments that measure the shuffling dynamics of KaiC-CI-S48T hexamers.

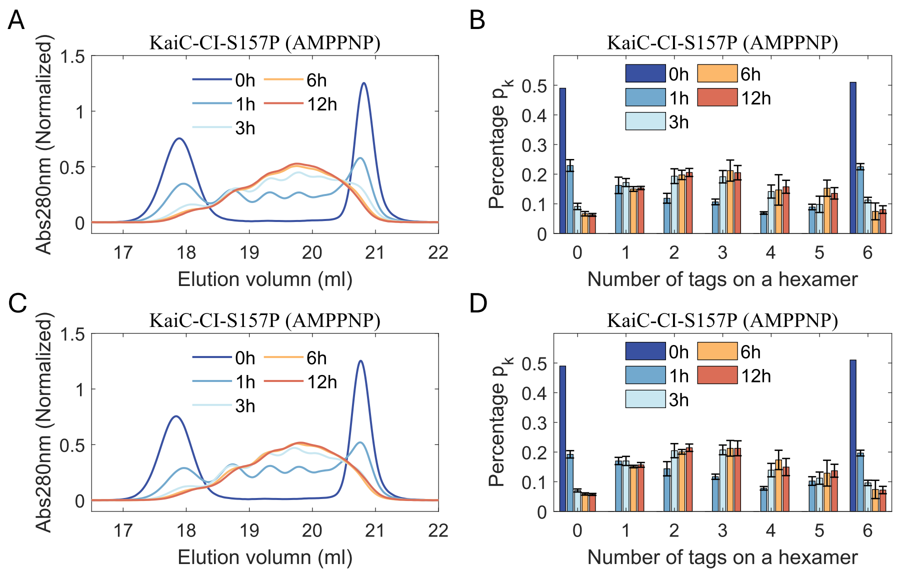


**Fig. S6.** Chromotography signals and recovered distributions for two independent experiments that measure the shuffling dynamics of KaiC-CI-S157P hexamers in reaction buffer where ATP is substituted by AMP-PNP.

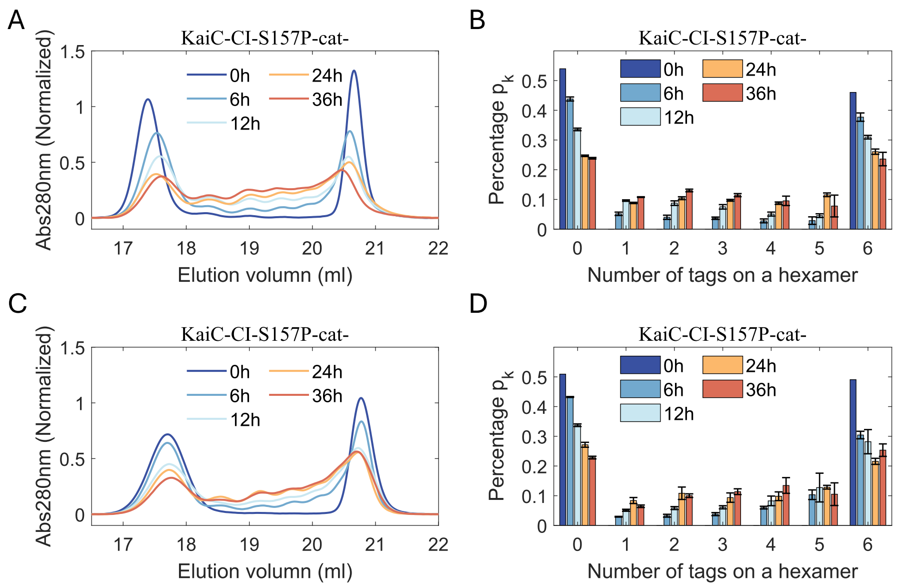


**Fig. S7.** Chromotography signals and recovered distributions for two independent experiments that measure the shuffling dynamics of KaiC-CI-S157P-E77Q/E78Q hexamers. E77 and E78 are catalytic sites for ATP of CI hexamers.

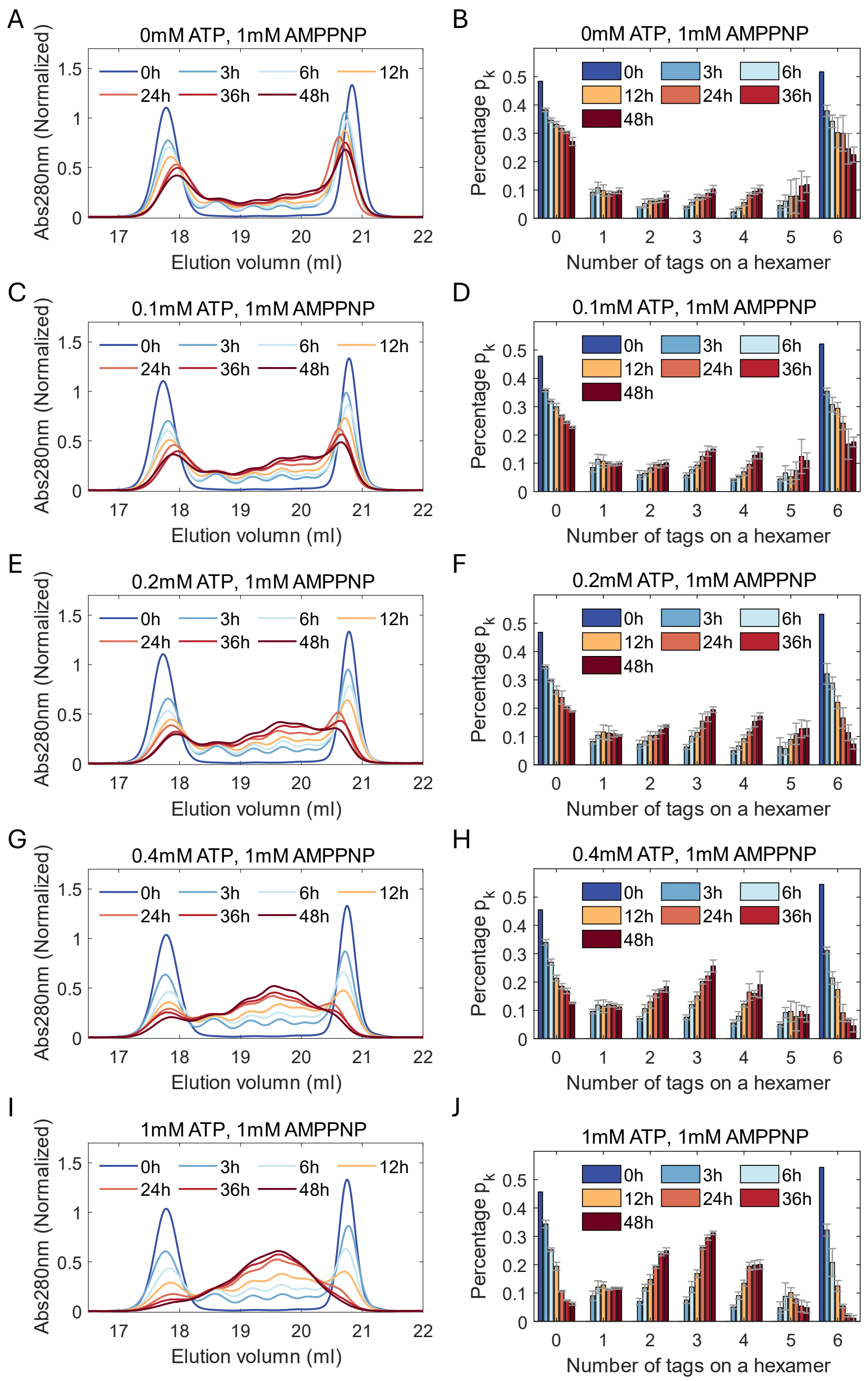


**Fig. S8.** Chromotography signals and recovered distributions for the first independent experiments (three in total) that measure the shuffling dynamics of KaiC-CI in reaction buffer with various concentrations of ATP and 1mM AMP-PNP.

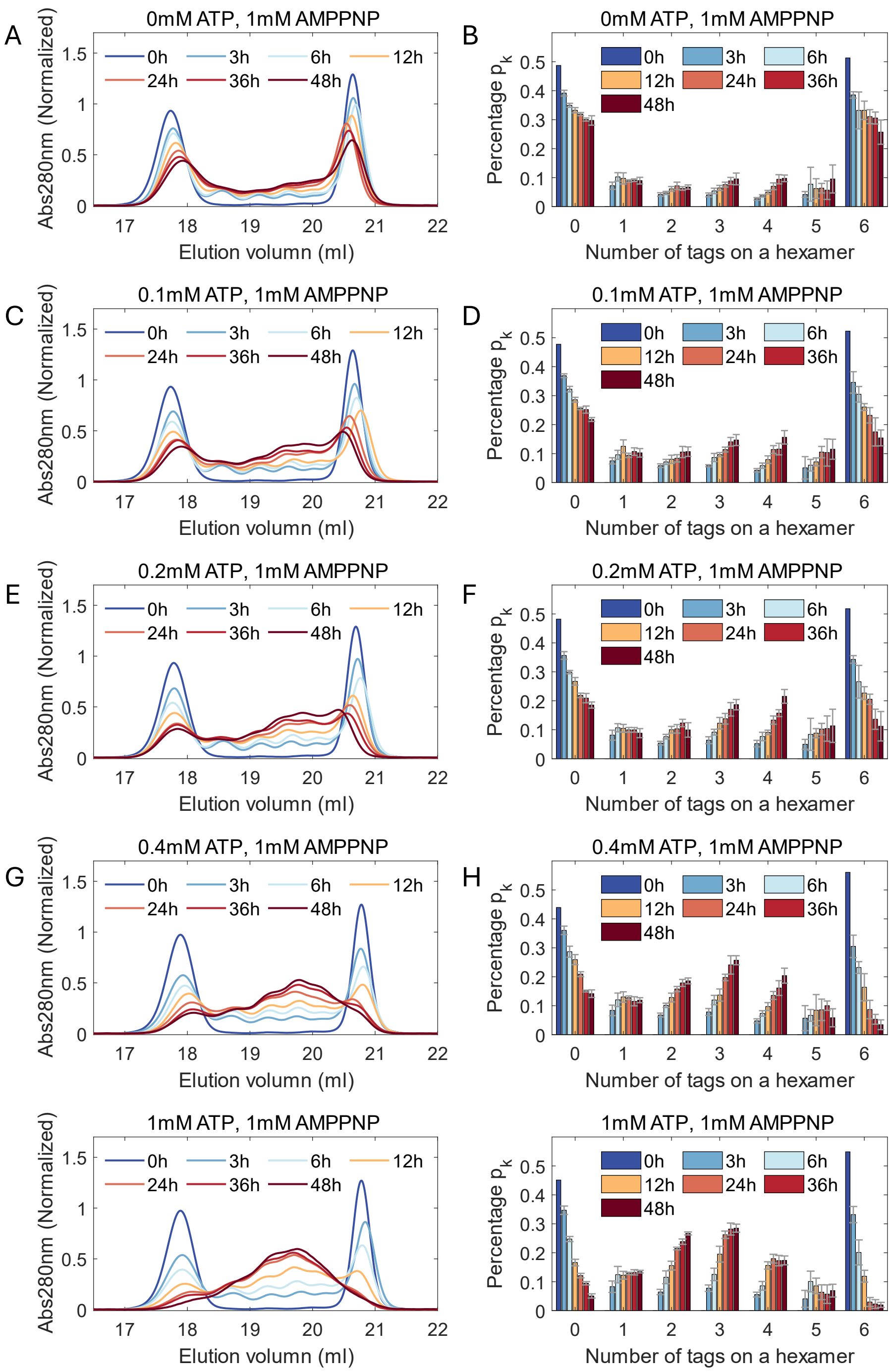


**Fig. S9.** Chromotography signals and recovered distributions for the second independent experiments (three in total) that measure the shuffling dynamics of KaiC-CI in reaction buffer with various concentrations of ATP and 1mM AMP-PNP.

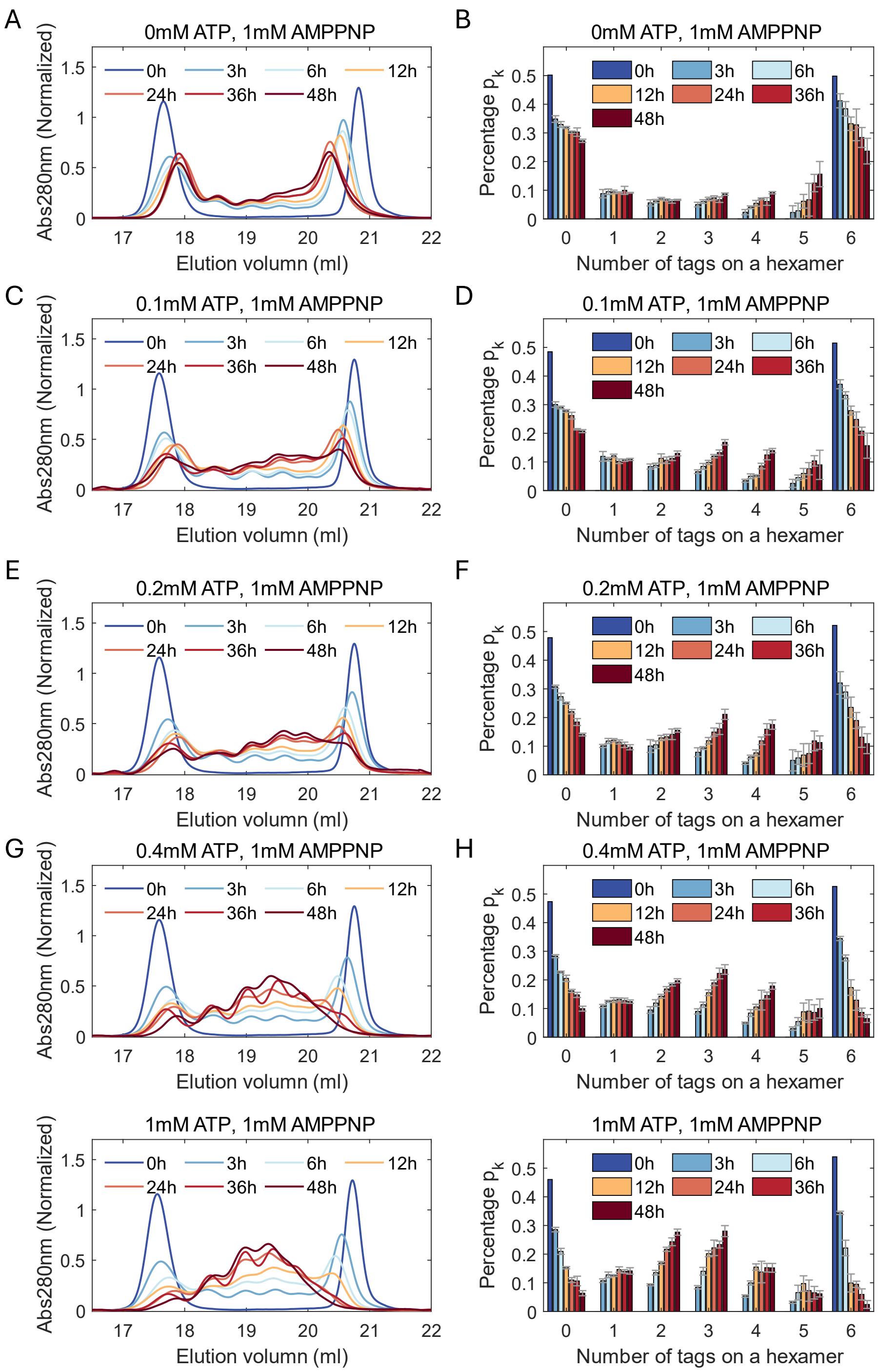


**Fig. S10.** Chromotography signals and recovered distributions for the third independent experiments (three in total) that measure the shuffling dynamics of KaiC-CI in reaction buffer with various concentrations of ATP and 1mM AMP-PNP.

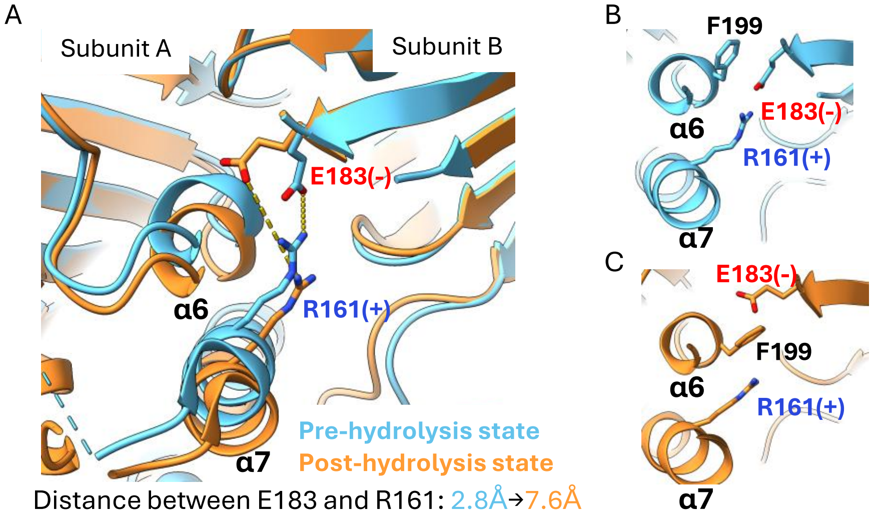


**Fig. S11.** Comparing the subunit-subunit interface of KaiC-CI hexamer. (A) Crystal structures of the pre and post-hydrolysis states are illustrated in the ribbon style[6]. After hydrolysis, besides the reposition of helix α6 and α7, a salt bridge between residue E183 and residue R161 is broken. (B-C) Local inspection of the salt bridge. In the post-hydrolysis state, repositioning of helix α6 induces the residue F199 to insert between E183 and R161 to further isolate their interactions, resulting in a loosened interface. Figures are generated with ChimeraX [7].

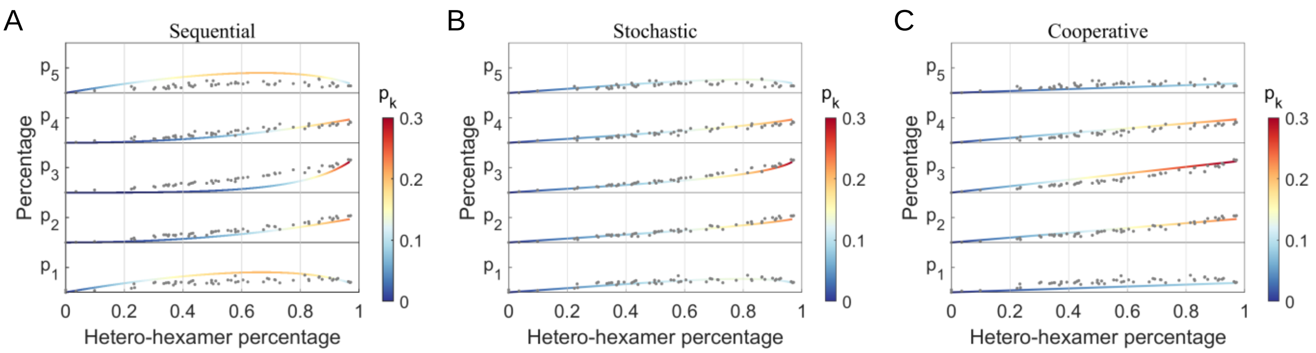


**Fig. S12.** Evolution of hetero-hexamer distribution to hetero-hexamer percentage for three dissociation patterns. Gray dots are experimental data (Fig. S1 and S3-S10).

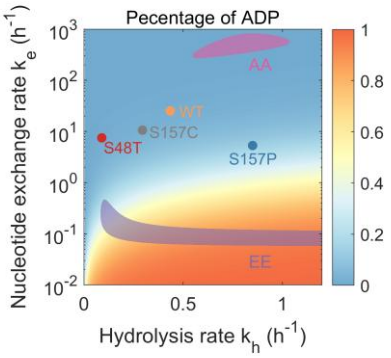


**Fig. S13** Prediction for percentage of ADP bound in CI domain.

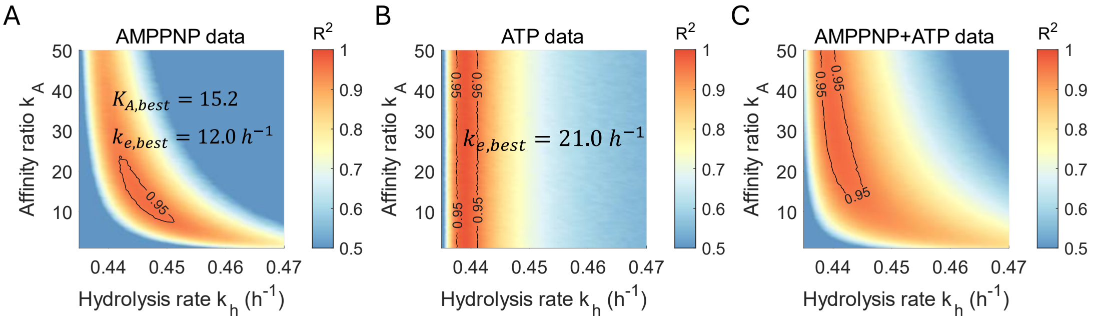


**Fig. S14.** Fitting for KaiC-CI-WT experimental data by directly stochastic simulation. Samples inhibited by AMP-PNP and incubated in 5mM ATP are fitted respectively and simultaneously. The region with $R^2$>0.95 is highlighted with black line.

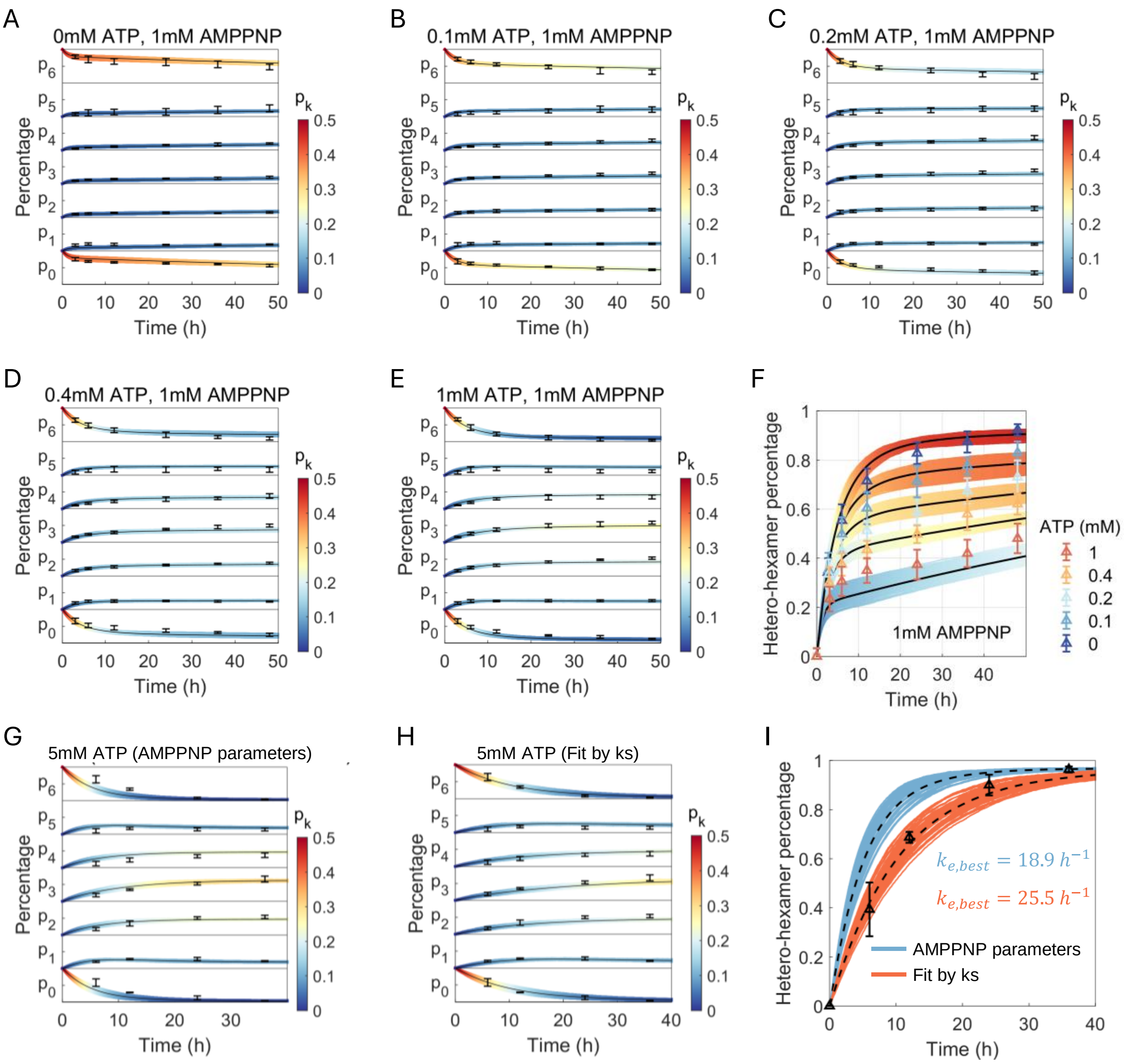


**Fig. S15** Fitting results of the experimental data. (A-F) Fitting results for KaiC-CI-WT incubating in buffer with 1mM AMP-PNP and various concentrations of ATP. The curve's width represents the fitting range (see method), and markers represent experimental data. (G-H) Fitting results for KaiC-CI-WT incubating in buffer with 5mM ATP.

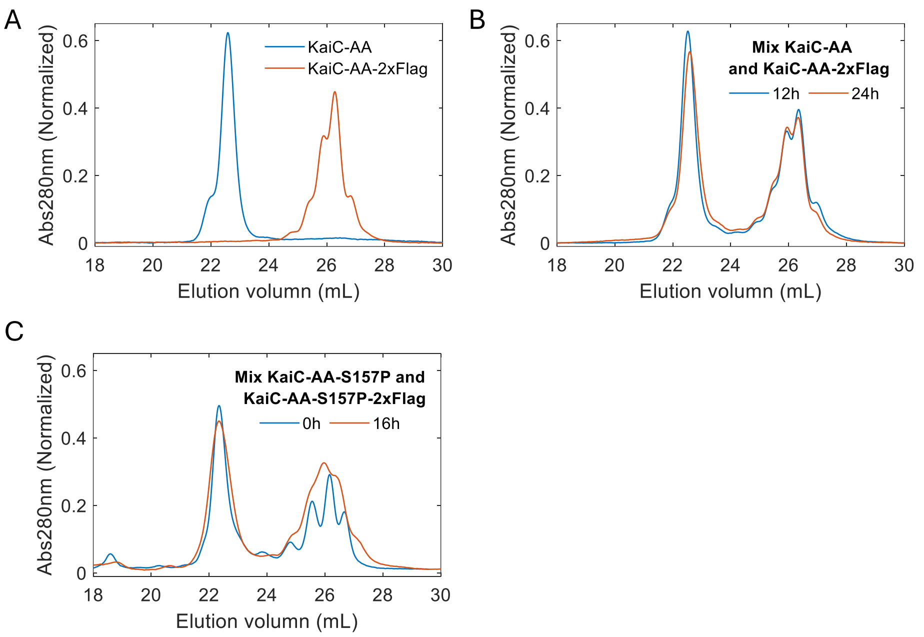


**Fig. S16** Chromatography signal for the shuffling dynamics of KaiC-AA hexamers. (A) Measurement for KaiC-AA and KaiC-AA-2×FLAG hexamers, respectively. KaiC-AA hexamers are eluted around 22.5ml, and tagged hexamers are eluted around 26ml. (B) Measurements after mixing KaiC-AA and KaiC-AA-2×FLAG hexamers in reaction buffer at 30°C in indicated time points. After incubating for 24 hours, the signal is just the combination of two signals in (A), suggesting that KaiC-AA hexamers are difficult to shuffle. (C) Messurments for KaiC-AA with S157P mutation. After incubating for 16 hours, the signal is still similar to the initial signal, indicating failure to shuffle.

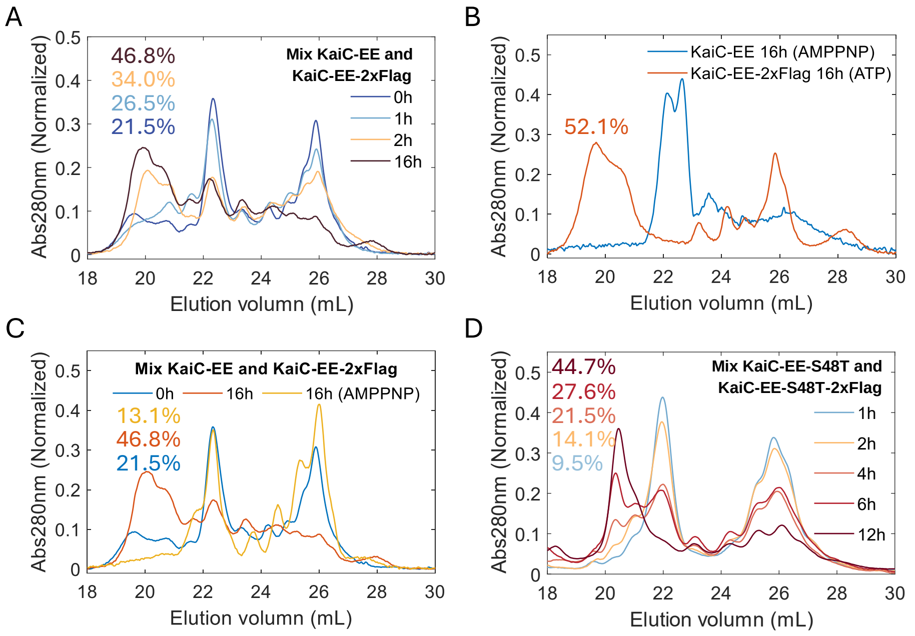


**Fig. S17** Chromatography signal for the shuffling dynamics of KaiC-EE hexamers. (A) Measurements after mixing KaiC-EE and KaiC-EE-2×FLAG hexamers in reaction buffer at 30°C in indicated time points. Initially, two peaks lie around 22.5ml and 26ml, respectively, as KaiC-AA in (Fig. S16 *A*). As time increases, no peaks emerge in the range corresponding to the hetero-hexamers, but intensity around 20ml increases. This portion of the sample contains both the native and tagged proteins, which are validated by SDS-PAGE(not shown). We assign this portion of protein to subunits that dissociated from a part of CS-hexamers, which were reported in a previous biochemical study[5]. The value indicated in the graph corresponds to this portion of signal. (B-C) Inhibiting formation of CS-hexamers and shuffling by replacing ATP with AMP-PNP in reaction buffer. After incubating for 16 hours, the intensity around 20ml is lower when inhibited by AMP-PNP. (D) Messurments for KaiC-EE with S48T mutation. As time increases, observable intensity increases around 20ml, indicating KaiC-EE-S48T can shuffle.

**Table S1 Model parameters fit by shuffling rate**

| Protein | ATPase activity* $k_{cat}$ $(h^{-1})$ | Shuffling rate $k_s$ $(h^{-1})$ | Nucleotide exchange rate $k_e$ $(h^{-1})$ | Hydrolysis rate $k_h$ $(h^{-1})$ |
|---|---|---|---|---|
| KaiC-CI-WT | 0.43 | 0.091±0.011 | 15.29~37.6 | 0.34~0.53 |
| KaiC-CI-S157P | 0.73 | 0.881±0.030 | 2.48~9.79 | 0.70~1.00 |
| KaiC-CI-S157C | 0.43 | 0.101±0.008 | 6.13~17.31 | 0.23~0.36 |
| KaiC-CI-S48T | 0.09 | 0.015±0.004 | 4.65~11.29 | 0.07~0.11 |

* ATPase activity is adapted from [6]
